# Anti-nucleolin aptamer, iSN04, inhibits the inflammatory responses in myoblasts by modulating the β-catenin/NF-κB signaling pathway

**DOI:** 10.1101/2023.04.01.535227

**Authors:** Machi Yamamoto, Mana Miyoshi, Kamino Morioka, Takakazu Mitani, Tomohide Takaya

**Affiliations:** Department of Agriculture, Graduate School of Science and Technology, Shinshu University, Nagano, Japan; Department of Agricultural and Life Sciences, Faculty of Agriculture, Shinshu University, Nagano, Japan; Department of Biomolecular Innovation, Institute for Biomedical Sciences, Shinshu University, Nagano, Japan

**Author notes:** Corresponding author: Tomohide Takaya E-mail address Department of Agricultural and Life Sciences, Faculty of Agriculture, Shinshu University, 8304 Minami-minowa, Kami-ina, Nagano 399-4598, Japan.

**Keywords:** β-catenin, inflammatory response, myoblast, myogenetic oligodeoxynucleotide (myoDN), nuclear factor-κB (NF-κB), nucleolin

## Abstract

A myogenetic oligodeoxynucleotide, iSN04, is the 18-base single-stranded DNA that acts as an anti-nucleolin aptamer. iSN04 has been reported to restore myogenic differentiation by suppressing inflammatory responses in myoblasts isolated from patients with diabetes or healthy myoblasts exposed to cancer-releasing factors. Thus, iSN04 is expected to be a nucleic acid drug for the muscle wasting associated with chronic diseases. The present study investigated the anti-inflammatory mechanism of iSN04 in the murine myoblast cell line C2C12. Tumor necrosis factor-α (TNF-α) or Toll-like receptor (TLR) ligands (Pam_3_CSK_4_ and FSL-1) induced nuclear translocation and transcriptional activity of nuclear factor-κB (NF-κB), resulting in upregulated expression of TNF-α and interleukin-6. Pre-treatment with iSN04 significantly suppressed these inflammatory responses by inhibiting the nuclear accumulation of β-catenin induced by TNF-α or TLR ligands. These results demonstrate that antagonizing nucleolin with iSN04 downregulates the inflammatory effect mediated by the β-catenin/NF-κB signaling pathway in myoblasts. In addition, the anti-inflammatory effects of iSN04 were also observed in smooth muscle cells and pre-adipocytes, suggesting that iSN04 may be useful in preventing inflammation induced by metabolic disorders.

## 1. Introduction

Skeletal muscle is the largest organ in humans (∼40% of body weight) [1], serves as the primary thermogenic tissue [2], stores 500 g of glycogen as an energy source [3], and absorbs approximately 80% of blood glucose in response to insulin [4]. Thus, muscle wasting disrupts systemic homeostasis and increases mortality in chronic diseases. Cancer cachexia is a complex syndrome involving irreversible muscle wasting [5] that occurs in 30-80% of cancer patients and accounts for about 20% of cancer deaths [6]. Muscle loss is also associated with diabetes mellitus [7] and is an independent risk for all-cause mortality in patients with type 2 diabetes [8]. Therefore, preventing disease-related muscle loss is an important therapeutic issue in clinical practice.

One of the causes of muscle wasting in chronic diseases is systemic inflammation induced by circulating factors such as pro-inflammatory cytokines and damage-associated molecular patterns (DAMPs) [9,10]. Cytokines such as interleukins (ILs) and tumor necrosis factor-α (TNF-α) activate the nuclear factor-κB (NF-κB) pathway, leading to inflammation, proteolysis, and dysregulation of myogenesis in skeletal muscle [11]. DAMPs are the diverse endogenous factors (nucleic acids, proteins, and proteoglycans) released from damaged organelles, dead cells, or disrupted tissues in various diseases [12]. Several DAMPs are recognized by Toll-like receptors (TLRs), initiate innate immune responses involving NF-κB activation, and eventually progress to muscle wasting [12,13]. Such inflammatory conditions and environments have been reported to disrupt myogenesis, which is driven by myogenic progenitor cells called myoblasts. For example, conditioned media from cancer cells strongly inhibit myoblast differentiation and myotube formation [14–17], or myoblasts isolated from patients with type 2 diabetes show impaired differentiation and autophagy [18,19]. These studies suggest that correcting myoblast function may have therapeutic benefits in inflammatory muscle wasting associated with chronic disease.

We recently identified the 18-base single-stranded DNA, iSN04, as a myogenetic oligodeoxynucleotide (myoDN) that acts as an anti-nucleolin aptamer and promotes myoblast and rhabdomyosarcoma differentiation [20–22]. iSN04 is taken up into the cytoplasm without a carrier and removes the translational inhibition of p53 mRNA by nucleolin, leading to activation of the p53 signaling pathway and the myogenic program [20]. iSN04 reverses the deteriorated differentiation of myoblasts isolated from patients with diabetes [23] and of healthy myoblasts exposed to cancer-conditioned media [24]. It shows that iSN04 can be a nucleic acid drug for muscle wasting by restoring myogenesis that has been impaired by diabetes or cancer. Interestingly, in addition to its myogenic ability, iSN04 has an anti-inflammatory effect. iSN04 suppresses the expression of IL-1β, IL-6, IL-8, and TNF-α induced by diabetes, high glucose, excess fatty acids, cancer-released factors, or TNF-α [23,24]. As these inflammatory responses are known to be mediated by NF-κB to disrupt myoblast differentiation [25], we hypothesized that the myogenic and anti-inflammatory effects of iSN04 are linked though nucleolin inhibition. A recent study reported that another anti-nucleolin aptamer, AS1411, suppressed NF-κB-regulated IL-6 expression in cardiomyocytes [26]. However, the anti-inflammatory mechanisms of anti-nucleolin aptamers remain unclear. This study investigated the effect of iSN04 on myoblast inflammation induced by TNF-α and TLR ligands, mimicking chronic diseases associated with systemic inflammation and skeletal muscle wasting.

## 2. Materials and Methods

### 2.1. Chemicals

Phosphorothioated iSN04 (5’-AGA TTA GGG TGA GGG TGA-3’) was synthesized and HPLC-purified (GeneDesign, Osaka, Japan) [20-24,27] and then dissolved in endotoxin-free water. Recombinant murine TNF-α (Fujifilm Wako Chemicals, Osaka, Japan), triacylated lipopeptide Pam_3_CSK_4_ (tripamitoyl-*S*-(bis(palmitoyloxy)propyl)-Cys-Ser-(Lys)_3_-Lys; Novus Biologicals, Centennial, CO, USA), and fibroblast-stimulating lipopeptide 1 (FSL-1) (*S*-(2,3-bispalmitoyloxypropyl)-Cys-Gly-Asp-Pro-Lys-His-Pro-Lys-Ser-Phe; Adipogen, San Diego, CA, USA) were rehydrated in phosphate-buffered saline. Equal volumes of the solvents were used as the negative controls.

### 2.2. Cell culture

Murine myoblast cell line C2C12 (DS Pharma Biomedical, Osaka, Japan) [20,23], rat embryonic thoracic aorta smooth muscle cell line A10 (CRL-1476; ATCC, Manassas, VA, USA) [28,29], and murine 3T3-L1 fibroblast cell line (IFO50416; JCRB Cell Bank, Osaka, Japan) as a model of pre-adipocytes [30] were cultured in DMEM (Nacalai, Osaka, Japan) supplemented with 10% fetal bovine serum (HyClone; GE Healthcare, Salt Lake City, UT, USA) and a mixture of 100 units/ml penicillin and 100 μg/ml streptomycin (Nacalai) at 37°C under 5% CO_2_ throughout the experiments. C2C12 cells were seeded on collagen type I-C (Cellmatrix; Nitta Gelatin, Osaka, Japan) coated dishes or plates.

### 2.3. Quantitative real-time RT-PCR (qPCR)

C2C12 cells were seeded on 30-mm collagen-coated dishes (1.5×10^5^ cells/dish), pre-treated with 10 μM iSN04 for 3 h, then additionally stimulated with 50 ng/ml TNF-α, 100 ng/ml Pam_3_CSK_4_, or 100 ng/ml FSL-1 for 2 h. A10 cells were seeded on 60-mm dishes (2.0×10^5^ cells/dish), pre-treated with 10 μM iSN04 for 3 h, then stimulated with 50 ng/ml TNF-α for 4 h. 3T3-L1 cells were seeded on 24-well plates (4.0×10^4^ cells/well), pre-treated with 30 μM iSN04 for 3 h, then stimulated with 5 ng/ml TNF-α or 10 ng/ml Pam_3_CSK_4_ for 2 h. Total RNA of the cells was isolated using NucleoSpin RNA Plus (Macherey-Nagel, Düren, Germany) and reverse transcribed using ReverTra Ace qPCR RT Master Mix (TOYOBO, Osaka, Japan). qPCR was performed using GoTaq qPCR Master Mix (Promega, Madison, WI, USA) with the StepOne Real-Time PCR System (Thermo Fisher Scientific, Waltham, MA, USA). The amount of each transcript was normalized to that of the murine 3-monooxygenase/tryptophan 5-monooxygenase activation protein zeta gene (*Ywhaz*) [31] in C2C12 and 3T3-L1 cells or the rat ribosomal protein L19 gene (*Rpl19*) [32] in A10 cells. Results are presented as fold-changes. Primer sequences of murine TNF-α (*Tnf*) [33], murine NF-κB p65 subunit (*Rela*) [34], murine IL-6 [35], rat IL-6 (*Il6*) [36], rat IL-8 (*Cxcl8*) [37], and rat monocyte chemoattractant protein 1 (MCP-1) (*Ccl2*) [37] have been described previously.

### 2.4. Dual-luciferase assay

5×NF-κB_RE::RedF (#124530; Addgene, Watertown, MA, USA) is the reporter plasmid consisting of a red firefly luciferase driven by a minimal promoter containing five NF-κB-binding sites (NF-κB-Luc) [38]. pRL-SV40 (#E2231; Promega) and pRL-CMV (#E2261; Promega) are the plasmid vectors consisting of a *Renilla reniformis* luciferase driven by simian virus 40 enhancer and early promoter elements (SV40-Luc) and by cytomegalovirus enhancer and immediate/early promoter elements (CMV-Luc), respectively. C2C12 cells were seeded on 12-well collagen-coated plates (1.5×10^4^ cells/well) and A10 cells were seeded on 12-well plates (3.0×10^4^ cells/well). The next day, the plasmids were transfected into the cells using Opti-MEM (Thermo Fisher Scientific) and Viofectin Transfection Reagent (Viogene, Taipei, Taiwan) according to the manufacturer’s instructions. After 6 h of transfection, the transfection medium was replaced with culture medium containing 10 μM iSN04. After 3 h of iSN04 treatment, 3 ng/ml (A10 cells) or 50 ng/ml (C2C12 cells) TNF-α or 100 ng/ml Pam_3_CSK_4_ was added to induce inflammatory responses. After 40 h of stimulation, the cells were harvested and the activities of firefly and *Renilla* luciferase were measured in the same cell lysate using the Dual-Luciferase Reporter Assay System (Promega) and GloMax Luminometer (Promega). Relative NF-κB promoter activities were calculated as the ratio of NF-κB-Luc to SV40-Luc (C2C12 cells) or CMV-Luc (A10 cells) [30,39].

### 2.5. Immunocytochemistry

C2C12 cells were fixed with 2% paraformaldehyde, permeabilized with 0.2% Triton X-100 (Nacalai), and immunostained with rabbit polyclonal anti-NF-κB p65 subunit antibody (ab16502; Abcam, Cambridge, UK) or rabbit polyclonal anti-β-catenin antibody (#9562; Cell Signaling Technology, Danvers, MA, USA). Alexa Fluor 488-conjugated donkey polyclonal anti-rabbit IgG antibody (Jackson ImmunoResearch, West Grove, PA, USA) was used as secondary antibody. Nuclei were stained with DAPI (Nacalai). Fluorescence images were captured using EVOS FL Auto microscope (AMAFD1000; Thermo Fisher Scientific).

### 2.6. Western blotting

Soluble whole-cell lysates of C2C12 cells were prepared using lysis buffer consisting of 0.1 M Tris-HCl (pH 7.4), 75 mM NaCl, and 1% Triton X-100 with protease inhibitor cocktail (1 mM 4-(2-aminoethyl)benzenesulfonyl fluoride hydrochloride, 0.8 μM aprotinin, 15 μM E-64, 20 μM leupeptin hemisulfate monohydrate, 50 μM bestatin, and 10 μM pepstatin A) (Nacalai). The lysates were denatured with 50 mM Tris-HCl (pH 6.8), 10% glycerol, 2% sodium dodecyl sulfate at 95°C for 5 min. Protein samples were subjected to SDS-PAGE on a 10% polyacrylamide gel followed by Western blotting using an iBlot 2 Dry Blotting System (Thermo Fisher Scientific). Rabbit polyclonal anti-β-catenin antibody, and mouse monoclonal anti-glyceraldehyde 3-phosphate dehydrogenase (GAPDH) antibody (5A12; Fujifilm Wako Chemicals) were used as primary antibodies. Horseradish peroxidase (HRP)-conjugated goat anti-rabbit and anti-mouse IgG antibodies (Jackson ImmunoResearch) were used as secondary antibodies, respectively. HRP activity was detected using ECL Prime Western Blotting Detection Reagent (Cytiva, Tokyo, Japan) and ImageQuant LAS 500 (GE Healthcare). Levels of β-catenin proteins were normalized to that of GAPDH using ImageJ software (National Institutes of Health, Bethesda, MD, USA).

### 2.7. Statistical analysis

Results are presented as the mean ± standard error. Statistical comparisons among multiple groups were performed using Tukey-Kramer test after one-way analysis of variance. Statistical significance was set at *p* < 0.05.

## 3. Results

### 3.1. iSN04 inhibits NF-κB-dependent inflammatory gene expression

Inflammatory responses in C2C12 myoblasts were induced by TNF-α, Pam_3_CSK_4_, or FSL-1. Pam_3_CSK_4_ and FSL-1 are the ligands for TLR1/2 and TLR2/6 heterodimers, respectively. As these TLR ligands induce inflammatory gene expression in myoblasts [40], we used them as mimics of DAMPs. As shown in Fig. 1A, qPCR revealed that all three stimuli strongly increased the mRNA levels of TNF-α (*Tnf*) and IL-6 (*Il6*) genes. Importantly, these inflammatory gene transcriptions were significantly suppressed by pre-treatment with iSN04. Although the expression of TNF-α and IL-6 is regulated by NF-κB [41], the mRNA levels of the NF-κB p65 subunit (*Rela*) were not altered by TNF-α, Pam_3_CSK_4_, FSL-1, or iSN04. On the other hand, luciferase assays showed that the promoter activities driven by NF-κB were definitely upregulated by TNF-α or Pam_3_CSK_4_ (Fig. 1B). These upregulations were significantly suppressed by pre-treatment with iSN04, as observed in qPCR experiments.

**Fig. 1.**
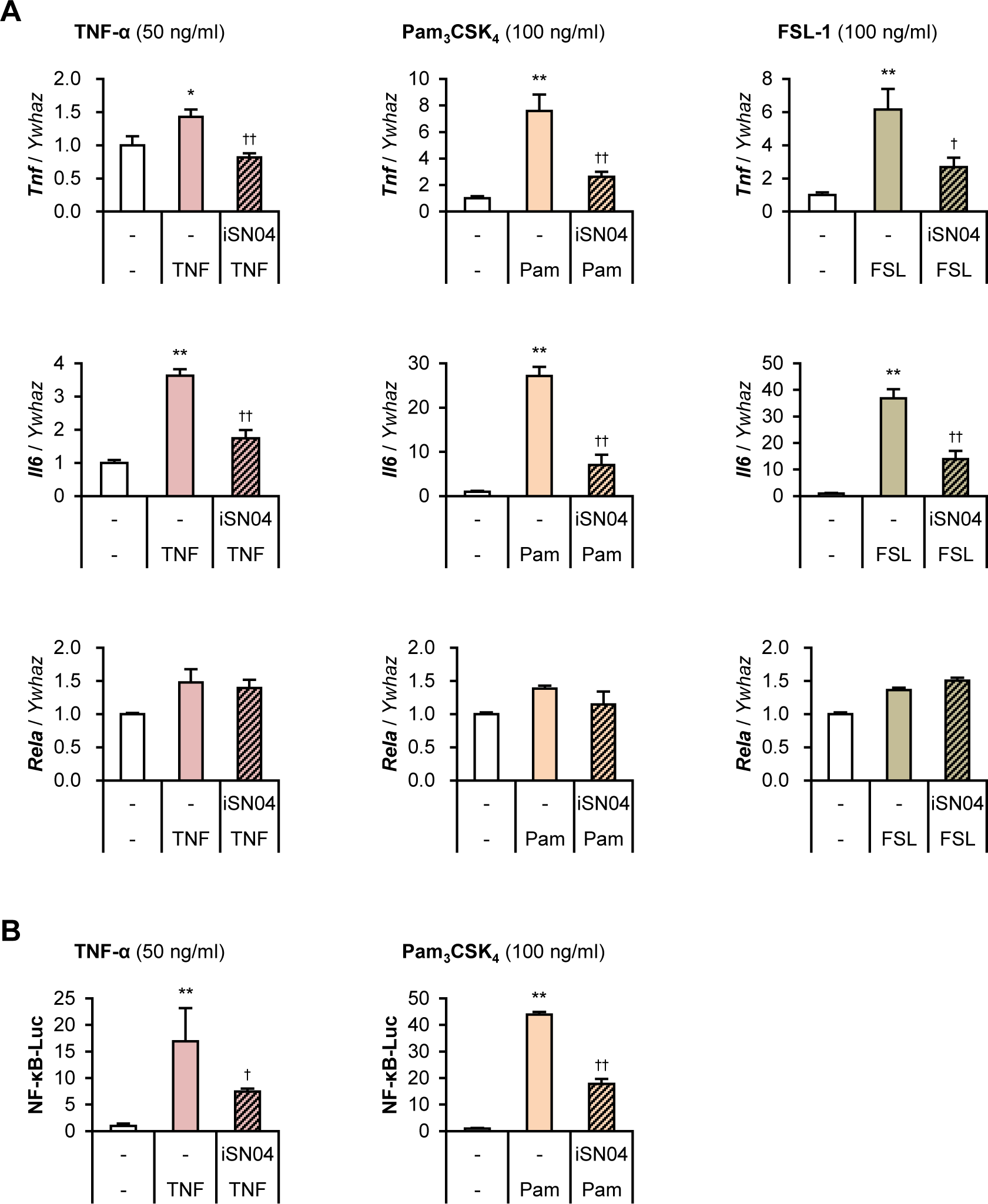
iSN04 inhibits NF-κB-dependent inflammatory gene expression in myoblasts. (A) qPCR results of TNF-α (*Tnf*), IL-6 (*Il6*), and NF-κB p65 subunit (*Rela*) expression in the C2C12 cells pre-treated with 10 μM iSN04 for 3 h and then treated with 50 ng/ml TNF-α, 100 ng/ml Pam_3_CSK_4_, or 100 ng/ml FSL-1 for 2 h. * *p* < 0.05, ** *p* < 0.01 vs control; ^†^ *p* < 0.05, ^††^ *p* < 0.05 vs ligand. *n* = 3-4. (B) Relative NF-κB-Luc activities in the C2C12 cells pre-treated with 10 μM iSN04 for 3 h and then treated with 50 ng/ml TNF-α or 100 ng/ml Pam_3_CSK_4_ for 40 h. ** *p* < 0.01 vs control; ^†^ *p* < 0.05, ^††^ *p* < 0.05 vs ligand. *n* = 3.

NF-κB transcribes inflammatory genes by translocating from the cytoplasm into the nucleus [41]. Immunostaining showed that TNF-α induced nuclear translocation of NF-κB in C2C12 cells within 30 min, as previously reported [42], but pre-treatment with iSN04 suppressed TNF-α-induced nuclear translocation of NF-κB (Fig. 2A). iSN04 similarly inhibited Pam_3_CSK_4_-induced nuclear translocation of NF-κB (Fig. 2B). These results demonstrate that iSN04 suppresses inflammatory gene expression by inhibiting NF-κB nuclear translocation in response to inflammatory stimuli.

**Fig. 2.**
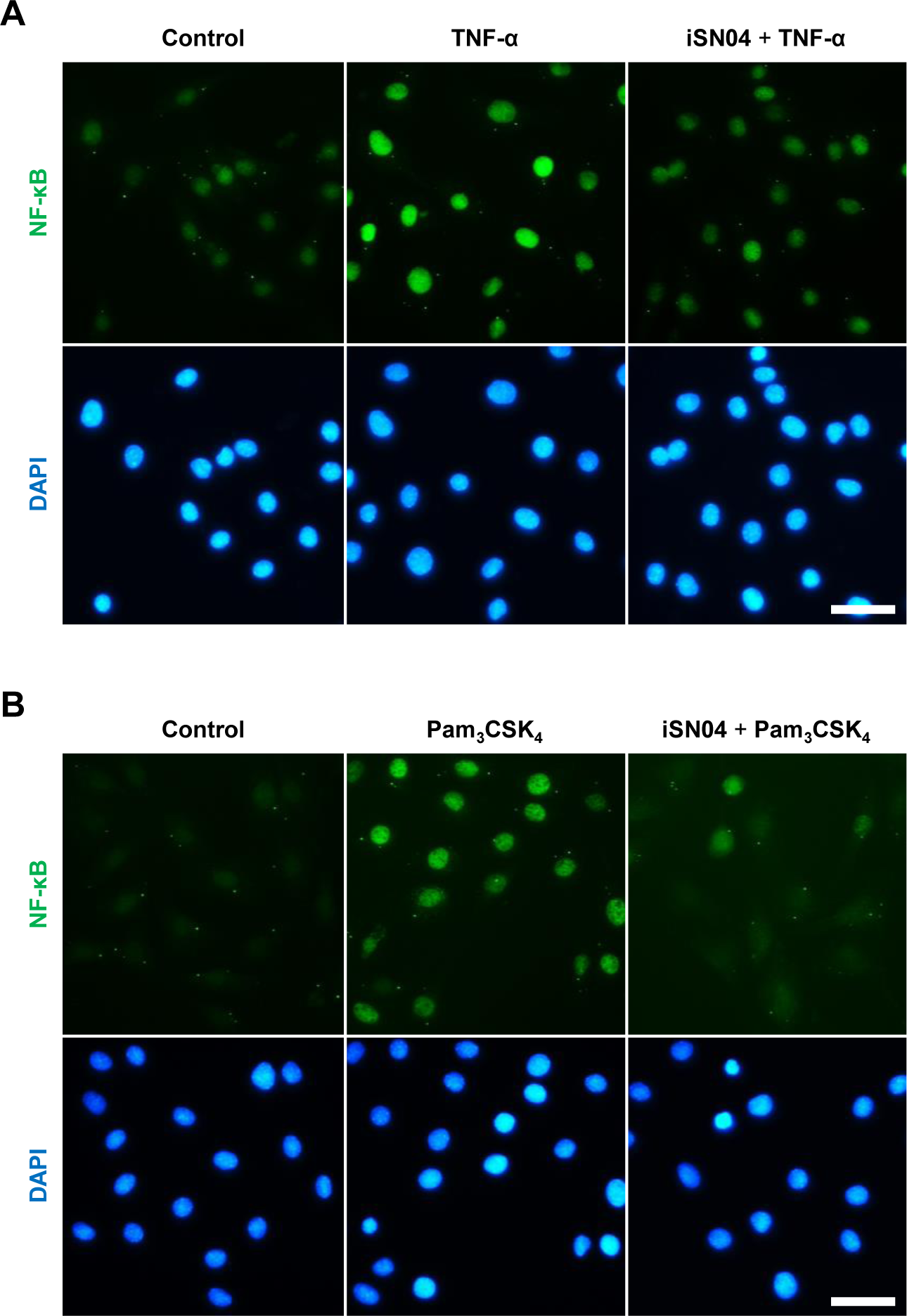
iSN04 inhibits nuclear translocation of NF-κB. (A and B) Representative images of NF-κB staining of the C2C12 cells pre-treated with 10 μM iSN04 for 3 h and then treated with 50 ng/ml TNF-α (A) or 100 ng/ml Pam_3_CSK_4_ (B) for 30 min. Scale bar, 50 μm.

### 3.2. iSN04 inhibits TNF-α-induced β-catenin accumulation

Nuclear translocation of NF-κB is mediated by β-catenin [43]. Immunostaining revealed the localization shift of β-catenin in C2C12 cells upon treatment with TNF-α and iSN04 (Fig. 3A). Consistent with previous studies [44,45], β-catenin localized to the cytoplasm under basal conditions and then translocated to the nucleus upon TNF-α stimulation. Interestingly, pre-treatment with iSN04 prevented TNF-α-induced nuclear translocation of β-catenin. Western blotting further demonstrated that the TNF-α-induced increase in β-catenin protein level was significantly suppressed by iSN04 treatment (Fig. 3B). These results suggest that iSN04 negatively regulates inflammatory responses by inhibiting the β-catenin/NF-κB signaling pathway in myoblasts.

**Fig. 3.**
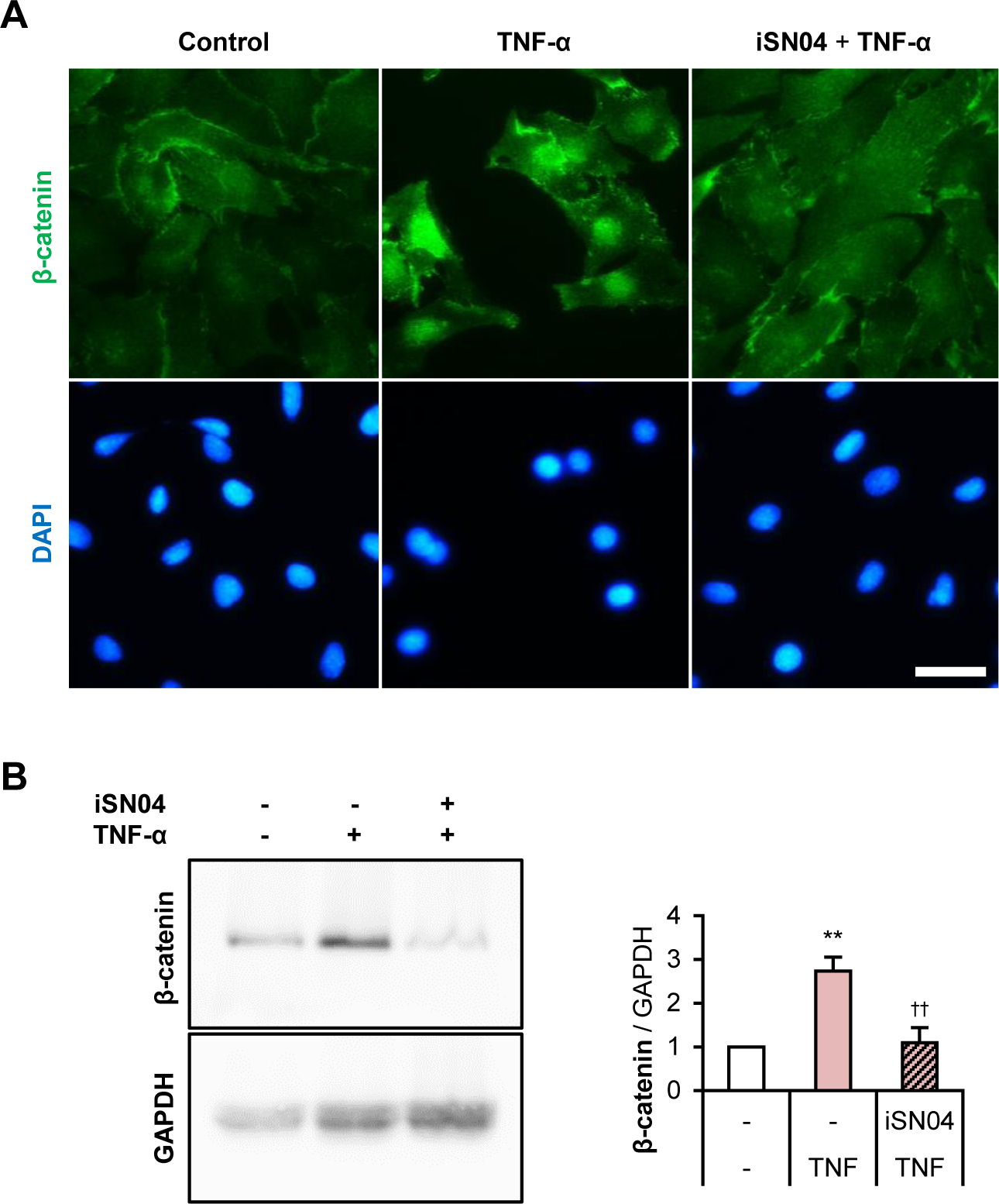
iSN04 inhibits GSK-3β phosphorylation and β-catenin activation. (A) Representative images of β-catenin staining of the C2C12 cells pre-treated with 10 μM iSN04 for 3 h and then treated with 50 ng/ml TNF-α for 1 h. Scale bar, 50 μm. (B) Representative images and quantification of Western blotting of β-catenin and GAPDH from C2C12 cells treated as in panel *A*. ** *p* < 0.01 vs control, ^†^ *p* < 0.05 vs TNF-α. *n* = 3.

### 3.3. iSN04 suppresses inflammation in smooth muscle cells and adipocytes

As nucleolin is ubiquitously expressed [45], antagonizing nucleolin may suppress inflammatory responses in different cell types, as reported in cardiomyocytes [26]. We investigated the effects of iSN04 on vascular smooth muscle cells (VSMCs) and pre-adipocytes, in which inflammation is a therapeutic target for atherosclerosis, diabetes, or obesity. Nucleolin has been reported to regulate the differentiation of VSMCs [28,29]. As shown in Fig. 4A, TNF-α significantly increased the mRNA levels of IL-6, IL-8 (*Cxcl8*), and MCP-1 (*Ccl2*) in rat VSMC line A10, but these inductions were significantly suppressed by pre-treatment with iSN04. Luciferase assay revealed that the TNF-α-induced transcriptional activity of NF-κB in A10 cells was significantly attenuated by iSN04 (Fig. 4B).

**Fig. 4.**
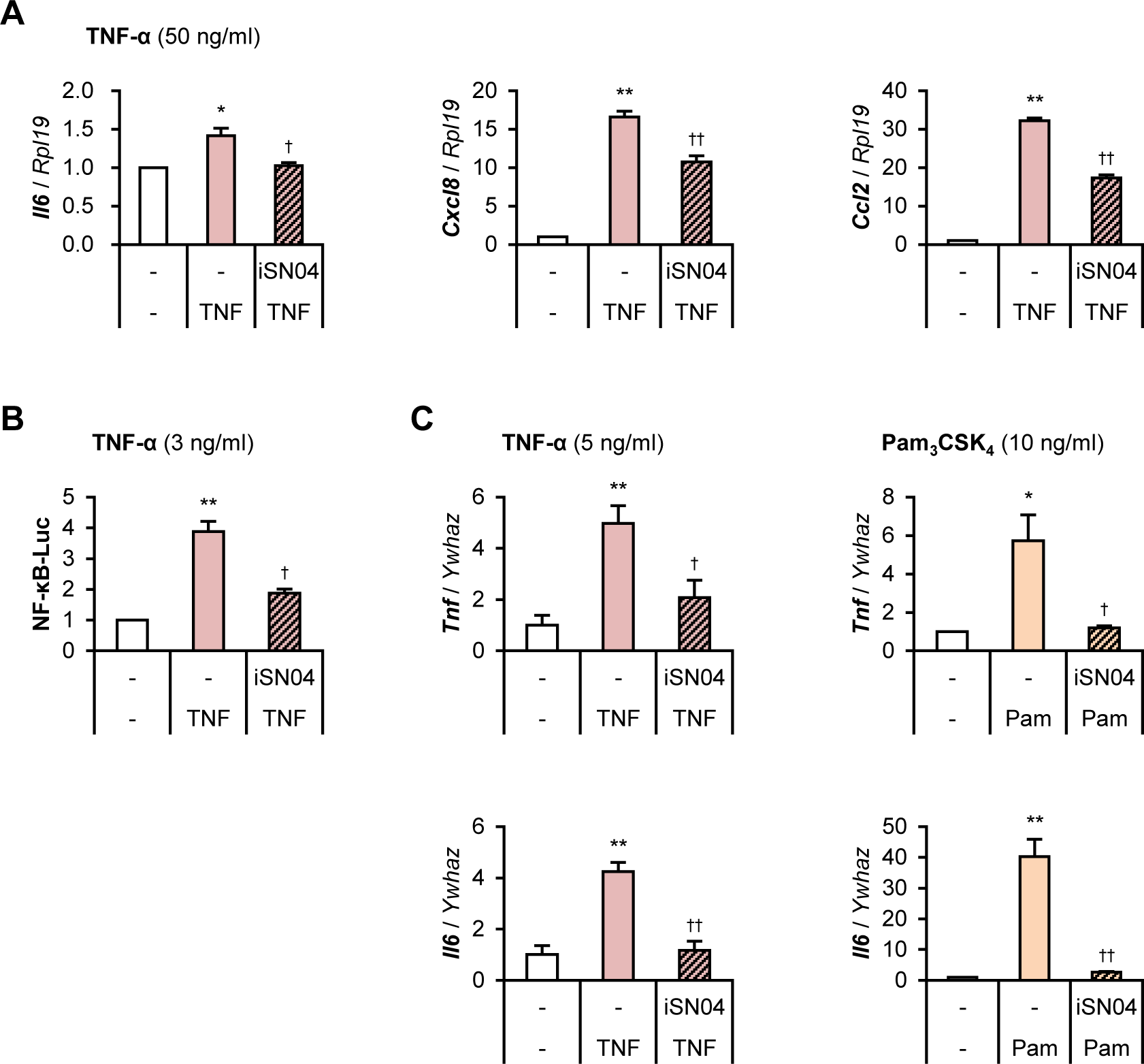
iSN04 inhibits NF-κB-dependent inflammatory gene expression in VSMCs and pre-adipocytes. (A) qPCR results of IL-6 (*Il6*), IL-8 (*Cxcl8*), and MCP-1 (*Ccl2*) expression in A10 cells pre-treated with 10 μM iSN04 for 3 h and subsequently treated with 50 ng/ml TNF-α for 4 h. * *p* < 0.05, ** *p* < 0.01 vs control; ^†^ *p* < 0.05, ^††^ *p* < 0.05 vs TNF-α. *n* = 3. (B) Relative NF-κB-Luc activities in the A10 cells pre-treated with 10 μM iSN04 for 3 h and subsequently treated with 3 ng/ml TNF-α for 40 h. ** *p* < 0.01 vs control, ^†^ *p* < 0.05 vs TNF-α. *n* = 3. (C) qPCR results of TNF-α (*Tnf*) and IL-6 (*Il6*) expression in the 3T3-L1 cells pre-treated with 30 μM iSN04 for 3 h and subsequently treated with 5 ng/ml TNF-α or 10 ng/ml Pam_3_CSK_4_ for 2 h. * *p* < 0.05, ** *p* < 0.01 vs control; ^†^ *p* < 0.05, ^††^ *p* < 0.05 vs ligand. *n* = 3-4.

In adipocytes, insulin induces nucleolin phosphorylation to regulate RNA efflux [47] and nucleolin binds to the leptin promoter as a transcriptional repressor [48]. The murine fibroblast cell line 3T3-L1 was used as a pre-adipocyte model to investigate the effect of iSN04. As shown in Fig. 4C, although both TNF-α and Pam_3_CSK_4_ induced the expression of TNF-α and IL-6, these inflammatory responses were significantly inhibited by pre-treatment with iSN04. These results observed in VSMCs and pre-adipocytes corresponded well to those observed in myoblasts, demonstrating that nucleolin antagonism by iSN04 exerts anti-inflammatory effects in multiple cell types.

## 4. Discussion

The present study showed that an anti-nucleolin aptamer, iSN04, suppressed inflammatory gene expression induced by TNF-α and TLR ligands in myoblasts, VSMCs, and pre-adipocytes. It is consistent with the previous report showing that another anti-nucleolin aptamer, AS1411, did so in cardiomyocytes [26]. By mechanism, iSN04 inhibited the nuclear translocation of NF-κB and reduced the promoter activity driven by NF-κB. In addition, iSN04 suppressed TNF-α-induced nuclear accumulation of β-catenin. It has been known that β-catenin interacts with NF-κB to form a transcriptional complex on their target genes in response to inflammatory stimuli [43]. Normally, the amount of free β-catenin in the cytosol is low, because β-catenin is phosphorylated by glycogen synthase kinase 3β (GSK-3β) and degraded by the proteasome [49]. GSK-3β also phosphorylates NF-κB to negatively regulate its activity [43]. Consequently, GSK-3β suppresses inflammatory gene expression mediated by the β-catenin/NF-κB signaling pathway. GSK-3β is inactivated by phosphorylation at Ser9 in a variety of biological processes [49]. It has been reported that nucleolin enhances GSK-3β phosphorylation and β-catenin-regulated transcription. Accordingly, nucleolin knockdown inhibits GSK-3β phosphorylation and reduces β-catenin protein [50]. A recent study showed that anti-nucleolin neutralizing antibody inhibits NF-κB nuclear translocation and inflammatory gene expression induced by lipopolysaccharide, a TLR4 ligand [51]. These findings suggest that nucleolin antagonism by iSN04 inhibits GSK-3β phosphorylation, which leads to β-catenin degradation and NF-κB inactivation, eventually exerting an anti-inflammatory effect.

In skeletal muscle, β-catenin/NF-κB signaling pathway dysregulates myogenesis [11], which has been implicated in inflammation-associated muscle wasting. As we have previously reported, iSN04 improves myogenic differentiation not only in normal conditions [20, 21], but also in diabetic and cachectic situations [23,24]. One of the mechanisms of iSN04-induced myogenesis has been hypothesized to be that inhibition of nucleolin increases p53 protein and activates its downstream [20]. The present study further suggests that suppression of the β-catenin/NF-κB signaling pathway by nucleolin antagonism may contribute to iSN04-induced myogenic differentiation. The dual action of iSN04 both in myogenesis and inflammation will have beneficial effects on muscle wasting associated with chronic inflammation in systemic diseases such as diabetes mellitus and cancer cachexia.

## Author contributions

TT designed the study and wrote the manuscript. MY, MM, KM, and TM performed experiments and analyses.

## Acknowledgments

This study was supported in part by grants from The Japan Society for the Promotion of Science to TT (19K05948 and 22K05554), Yamaguchi Educational and Scholarship Foundation to TT, and The Fund of Nagano Prefecture to Promote Scientific Activity to MY, MM, and KM (NPS2022325, NPS2021320, and NPS2021321).

